# Myeloid Specific Ablation of SHIP1 Boosts ex vivo Expansion and Regulatory Function of Myeloid-Derived Suppressor Cells in Inflammatory Arthritis

**DOI:** 10.1101/2025.09.30.679529

**Authors:** Eui Young So, Moon Jung Choi, Young Eun Lee, Bedia Akosman, Euy-Myoung Jeong, Anthony M. Reginato, Olin D. Liang

**Affiliations:** Division of Hematology/Oncology, Rhode Island Hospital, Warren Alpert Medical School of Brown University, Providence, Rhode Island, USA; Division of Rheumatology, Department of Medicine, Rhode Island Hospital, Warren Alpert Medical School of Brown University, Providence, Rhode Island, USA

**Author notes:** Correspondence: Olin D. Liang, Ph.D., One Hoppin Street, CORO West Research Building 4th Floor Suite 4.200, Providence, Rhode Island 02903, USA. These authors contributed equally to this work (Eui Young So, Moon Jung Choi).

**Keywords:** lipid phosphatase SHIP1, myeloid-derived suppressor cells, collagen-induced inflammatory arthritis, rheumatoid arthritis

## Abstract

Myeloid-derived suppressor cells (MDSCs) are a heterogeneous cell population and the immunosuppressive function of MDSCs has been well established in tumor microenvironment. Recent studies show that adoptive transfer of MDSCs can ameliorate collagen-induced inflammatory arthritis (CIA), a mouse model of human rheumatoid arthritis (RA). Src homology 2 domain-containing inositol polyphosphate 5-phosphatase 1 (SHIP1) was previously shown to regulate MDSC differentiation. In this study, we aimed to generate immunosuppressive MDSCs from mouse bone marrow (BM) through genetic modification combined with cytokine treatments, and to investigate the ability of these ex vivo induced BM-MDSCs to suppress inflammatory responses in the CIA mouse model of RA. We found that myeloid specific ablation of SHIP1 increased the ratio of MDSCs and enhanced their regulatory functions in cytokine induced BM culture. MDSCs from LysMcre:SHIP1^flox/flox^ mouse BM culture demonstrated stronger inhibitory effect on T cell proliferation than those from control mouse BM. Ex vivo induced MDSCs from either control mice or mice with myeloid specific ablation of SHIP1 were administered to the CIA mice as a cell-based therapy to treat inflammatory arthritis. Adoptive transfer of either BM-MDSCs significantly reduced disease incidence and severity, but SHIP1 deficient BM-MDSCs exhibited even higher efficacy compared to wild-type BM-MDSCs. Furthermore, pharmacological inhibition of SHIP1 enhanced the expression of immune regulatory genes in BM-derived MDSCs, and adoptive transfer of these cells protected against CIA development. In conclusion, myeloid specific ablation of SHIP1 boosts ex vivo expansion and immune regulatory function of MDSCs in experimental inflammatory arthritis. These ex vivo generated BM-MDSCs may provide novel therapeutic opportunities for the treatment of RA and other inflammatory diseases.

## Introduction

Rheumatoid arthritis (RA) is an autoimmune disease characterized by infiltration and accumulation of autoreactive immune cells in synovial joints. Joint inflammation is associated with joint pain, swelling and stiffness, joint surface erosion and bone resorption, leading to joint loss of function and disability [1–3]. If the disease remains untreated, it can cause severe joint damage, bone destruction, poor quality of life, and high morbidity and mortality [4]. In RA, cells within the inflamed synovium proliferate and invade the bone, activating bone resorption by osteoclasts. CD4 T cell infiltration is a hallmark of the pathogenesis of RA, and the immunogenetics of RA suggest a key role for aberrant pathways of T cell activation in the initiation and/or perpetuation of disease [5]. The synovial tissue and synovial fluid home in many types of innate effector cells as well, including but not limited to macrophages, mast cells and natural killer cells. These cells secrete several cytokines and inflammatory mediators, including TNF-α, IL-1 and IL-6. This inflammatory microenvironment within the joint supports the differentiation of pro-inflammatory T cells (Th17) and inhibits the development of regulatory T cells (T_reg_). An imbalance between Th17 and T_reg_ has been associated with RA severity and subsequent joint destruction [6, 7].

Myeloid-Derived Suppressor cells (MDSCs) are a heterogeneous cell population that consist of myeloid progenitor (MP), immature dendritic (DCs)/granulocytes/macrophages cells. MDSCs are classified by surface markers (CD11b and Gr-1, in mouse) and functional markers, such as Arg-1 and iNOS. MDSCs can suppress immune response through depleting amino acids, including L-arginine and cysteine, by Arg-1 and iNOS [8]. Because these amino acids are critical for proliferation and function of T cells, infiltrated MDSCs suppress proliferation and activity of T cells in tumor or inflammation sites. MDSCs also suppress NK cell cytotoxicity by secretion of TGF-β [9], and IL-10 and IFN-γ secreted from MDSCs induce the activation of T_reg_ [10]. Although the immunosuppressive function of MDSCs has been well established in cancers [11, 12], their regulation in autoimmune diseases is not fully understood. Recent studies showed that adoptive transfer of MDSCs ameliorates collagen-induced inflammatory arthritis (CIA) [13, 14]. However, antigen dependent maturation, weak regulatory function of naïve MDSCs, and limited number of functional MDSCs in normal host make it difficult to use MDSCs as a therapeutic tool for treatment. Therefore, specific modification tools to enhance regulatory function and control proliferation and development of MDSCs should be considered for therapeutic application for inflammatory disease, including RA.

The SH2-containing inositol-5’-phosphatase-1 (SHIP1) controls PI3K signaling by limiting membrane recruitment and activation of AKT [15]. Systemic knock-out (KO) of the SHIP1 gene in mice leads to an increase in MDSCs as well as T_reg_ [16, 17], and abnormal early B cell development in mice [18]. However, genetic loss of SHIP1 also induces pathological condition mediated by increased cytokines and inflammation in mice [19–21]. Since earlier studies have shown that MDSCs and T_reg_ cells have immunosuppressive effects under disease conditions [22], cell modification and reprograming microenvironment by inhibition of SHIP1 could be a promising approach for autoimmune diseases, including RA. Therefore, our studies have focused on investigating whether targeting SHIP1 can change cellular repertoire or activation and suppress the development of inflammatory disease, such as RA. In our previous study, treatment of mouse bone marrow (BM) cells with 3AC led to an expansion of myeloid populations [23]. Using the CIA mouse model, mice treated with the SHIP1 inhibitor, 3α-amino cholestane (3AC) exhibited a marked expansion of MDSCs in vivo. Furthermore, we found that transplantation of MDSCs isolated from 3AC-treated mice into CIA mice alleviated the disease, suggesting that the MDSCs were the cellular mediators of the observed beneficial effects [24]. Because MDSCs have a remarkable ability to suppress the immune responses of pro-inflammatory Th17, ex vivo expanded MDSCs can then be administered intravenously back to the patient as an autologous cell-based therapy to treat RA and potentially other autoimmune diseases e.g., psoriatic arthritis [25] and systemic lupus erythematosus [26]. In the present study, we isolated BM mononuclear cells from mice with myeloid specific SHIP1 ablation (LysMcre:SHIP1^flox/flox^) and from SHIP1^flox/flox^ control mice and induced a substantial ex vivo expansion of the MDSCs in culture. Additionally, we used SHIP1-specific inhibitor to induce temporal SHIP1 loss in BM-MDSCs. We have also investigated the anti-inflammatory effect of these BM-MDSCs in the CIA mouse model of RA. Our study suggests that myeloid specific ablation of SHIP1 boosts ex vivo expansion and immune regulatory function of MDSCs in experimental inflammatory arthritis. These ex vivo generated BM-MDSCs may provide novel therapeutic opportunities for the treatment of RA and other inflammatory diseases.

## Materials and methods

### Mice

8∼10-week-old male DBA/1J (Strain # 000670) mice were purchased from Jackson Lab. SHIP1^flox/flox^ mice were a gift from Dr. William G. Kerr (Upstate Medical University, NY) and backcrossed into DBA/1J strain (F10) for experiments. LysMcre transgenic mice (B6.129P2- Lyz2^tm1(cre)Ifo^/J, Strain # 004781) were purchased from The Jackson Laboratory (Bar Harbor, ME) and backcrossed into DBA background (F10). The mice were then crossbred with SHIP1^flox/flox^ mice to generate mice with myeloid specific deletion of SHIP1 (LysMcre:SHIP1^flox/flox^). SHIP1^flox/flox^ mice were used as control. All mice were housed in a pathogen-free laboratory at the research institute of Brown University Health. All protocols of these experiments have been approved by the IACUC of Brown University Health.

### Flow cytometry analysis

Peripheral blood (PB) samples were collected from the retro-orbital sinus of mice; bone marrow (BM) of femurs and tibiae, and spleens were isolated from sacrificed mice. PB mononuclear cells (PBMCs), whole BM cells and splenocytes were resuspended into a single cell suspension in 10 ml ice cold phosphate-buffered saline (PBS) containing 2% fetal bovine serum (FBS). Cells were pelleted by centrifugation at 350g for 5 min, and supernatant was discarded. Following treatment with 3 ml of 1 x Red Blood Cell Lysis Buffer (Cat. # 420301, BioLegend, San Diego, CA) on ice for 10 min, cell lysis was stopped by adding 10 ml ice cold PBS containing 2% FBS. Cells were pelleted by centrifugation at 350g for 5 min, and supernatant was discarded. Cells were resuspended, filtered through a 35 µm nylon mesh and washed twice with 10 ml ice cold PBS containing 2% FBS, these cells were then prepared to a concentration of 10 x 10^6^ cells/ml in ice cold PBS containing 2% FBS, and 100 µl of cells were distributed into 5 ml plastic tubes. Samples were incubated with the TruStain FcX™ PLUS (Cat. # 156603, BioLegend) containing anti-mouse CD16/32 antibody at 0.25 µg/sample for 10 min on ice to block the Fc Receptors. Fluorophore-conjugated antibodies were then added to the cells (1 µl of each antibody per sample) and the cell suspensions were briefly vortexed and incubated on ice in the dark for 30 min. The incubation was stopped by adding 3 ml ice cold PBS containing 2% FBS. Cells were pelleted by centrifugation at 350g for 5 min, and supernatant was discarded. The cell pellets were resuspended in 0.5 ml of ice-cold PBS-FBS containing 4’,6- Diamidino-2-Phenylindole (DAPI, 1:1000 dilution, Cat. # 422801, BioLegend) to exclude DAPI+ dead cells during flow cytometry analysis. Labeled cells were immediately subjected to flow cytometry analysis. Except PE-anti-SHIP1 antibody (Santa Cruz, TX), all other antibodies used in flow cytometry were purchased from BioLegend (San Diego, CA): Alexa Fluor 488 anti-mouse CD11b, APC anti-mouse Gr-1, PE anti-mouse CD3e, APC/Cy7 anti-mouse B220, Alexa Flour 488 anti-mouse CD4, and APC anti-mouse CD25. Data were acquired using LSR II flow cytometer (BD Biosciences) and analyzed with FlowJo software (FlowJo, Ashland, OR).

### BM-MDSC culture

BM cells were obtained by flushing the femur and tibia of 5∼8 weeks old mice with 1 x HBSS. After lysis of red blood cells with RBC-lysis solution, cells were cultured in RPMI media supplemented with 10% FBS and penicillin/streptomycin/amphotericin B (Thermo Fisher, Waltham, MA) containing GM-CSF (10 ng/ml) or GM-CSF and IL-6 (10 ng/ml) (PeproTech, Cranbury, NJ) as described [27, 28]. To inhibit SHIP1 in BM-MDSC, 3-a-Aminocholestane (3AC, ECHELON Biosciences, UT) was treated at 1 µg/ml in BM-culture. To stop the maturation of myeloid cells and enrich MDSCs, suspended cells were harvested on Day 3∼5 and stained with antibodies against CD11b and Gr-1 and then analyzed by flow cytometer to evaluate the efficiency of MDSC differentiation.

### Generation of lentivirus and transduction of BM culture

To knock out SHIP1 gene expression in BM culture, Cre-containing plasmid [29] was cloned into a lentiviral shuttle vector and used for generation of virus. Lentivirus were produced by transfection of HEK293FT cells (Invitrogen) with 1 shuttle and 4 packing plasmids as described [30]. All plasmids for lentivirus production were purchased from Addgene (Watertown, MA). Culture media were collected at 48, 72 and 96 hours, cleared by low-speed centrifugation and filtration. All viruses were concentrated by ultracentrifugation at 23,200 rpm and titrated by the p24 ELISA Kit (R & D Systems, Minneapolis, MN). BM cells were isolated from SHIP1^flox/flox^ mice and infected with virus using the RetroNectin (Takara Bio, San Jose, CA)-bound method at Day 0 of GM-CSF-induced MDSC culture. Cells were harvested and subjected to flow cytometry and real-time PCR at Day 4.

### RNA preparation and real time PCR

Total RNA was isolated from BM-MDSCs using the TRIzol reagent (Invitrogen, Waltham, MA). First strand cDNA was synthesized by using the iScript Advanced cDNA Synthesis kit (BioRad, Hercules, CA) according to the manufacturer’s instructions. Primers for mouse genes encoding Arg1, iNos, and GAPDH were purchased from Qiagen. For quantitative analysis of gene expression, real-time PCR was performed using iTaq Universal Supermix in a CFX Connect real-time PCR detection system (Bio-Rad). Relative gene expression level was expressed by the fold increase over control sample. GAPDH gene was used as an internal control to validate the total transcript.

### In vitro T cell proliferation assay

To examine the regulatory function of MDSCs for T cells, mouse T cells were isolated from the spleen, using EasyStep mouse T cell isolation kit (STEMCELL Technologies). T cells were labeled with CFSE according to the manufacturer’s instructions (Biolegend). CFSE positive T cells were stimulated with Con A (eBioscience) and co-cultured at 1:1, 2:1 and 4:1 ratio with BM-MDSCs in 96-well plates. T cell proliferation was determined by intracellular intensity of CFSE using flow cytometry after 48 hours in culture.

### Collagen-induced inflammatory arthritis (CIA) and assessment of arthritis development

CIA was described previously [24]. Briefly, mice were injected into the base of the tail with a total of 100 µl complete Freund’s adjuvant (CFA) emulsion containing 100 µg Chicken Collagen II (Sigma-Aldrich, St. Louis, MO) at Day 0. Collagen II in incomplete Freund’s adjuvant (IFA) was injected on Day 21 as a second immunization. The development and severity of CIA in mice were assed using an established mouse scoring system for arthritis (30). For photographic and x-ray imaging, mice were euthanized at Day 50, and limbs were collected and fixed. After photographic analysis, fixed limbs were scanned with Faxitron cabinet X-ray system (Tucson, AZ).

### Adoptive transfer of MDSCs

After dual staining for CD11b and Gr-1, MDSCs were sorted from BM culture to obtain highly purified BM-MDSCs (>98%) using an Influx cell sorter (BD Biosciences). Isolated cells were transplanted into DBA mice through tail vein injection (1 x 10^6^ cells/mouse) on Day 0 and Day 20 before first and second immunization in the CIA mouse model of RA. DBA mice injected with PBS without MDSCs were used as controls.

### Statistical analysis

Analysis of statistical significance for all experiments was performed using the student’s t-test with GraphPad Prism software. P-values less than 0.05 were considered significant.

## Results

### Myeloid specific ablation of SHIP1 promotes the expansion of MDSCs

MDSCs are defined as Gr1+CD11b+ cells and have inhibitory function on development and activation of T cells [31]. Global knockout (KO) of SHIP1 in mice caused abnormal cellular development and led to apparent health complications [32], whereas LysMcre:SHIP1^flox/flox^ mice have a normal life span with no critical health problem [33]. Here, we used LysMcre:SHIP1^flox/flox^ mice and SHIP1^flox/flox^ control mice for BM isolation and downstream experiments. First, we examined whether myeloid restricted loss of SHIP1 would induce an expansion of the immunoregulatory cells in vivo. Analyses of splenocytes and peripheral blood mononuclear cells (PBMCs) suggested that the frequency of MDSCs increased in peripheral tissues of LysMcre:SHIP1^flox/flox^ mice compared to SHIP1^flox/flox^ control mice (Figures 1A, B). There was no difference in the frequency of MDSCs in BM between LysMcre:SHIP1^flox/flox^ and SHIP1^flox/flox^ mice (Fig1 C). We also observed significantly increased T_reg_ cell number in the spleen of LysMcre:SHIP1^flox/flox^ mice (Figure 1D), which is consistent with previous studies[34, 35]. However, expansion of the immunoregulatory compartment in LysMcre:SHIP1^flox/flox^ mice remained moderate in comparison with that in SHIP1 KO mice [36]. Next, we sought to examine the influence of myeloid expression of SHIP1 on the development and function of BM-derived MDSCs in vitro. BM-MDSCs were differentiated in BM cultures containing GM-CSF or GM-CSF + IL-6, and MDSCs were defined as CD11b+Gr1+ cells (Figure 2A). We found that either GM-CSF alone or GM-CSF with IL-6 co-stimulation could significantly increase both the cell number and frequency of MDSCs after 4-5 days of culture, which agrees with an earlier study published by Lee et al. [37]. We also found that SHIP1 is gradually expressed in GM-CSF-containing media in time-dependent manner (Supplementary Figure 1). More importantly, while there was no difference in total cell number, the frequency of BM-MDSCs from LysMcre:SHIP1^flox/flox^ BM cultures was significantly higher than that from the control SHIP1^flox/flox^ BM cultures, (Figures 2B, C). Hence, myeloid specific loss of SHIP1 suppresses cell maturation and increases the ratio of MDSCs in GM-CSF-induced BM culture.

**Figure 1.**
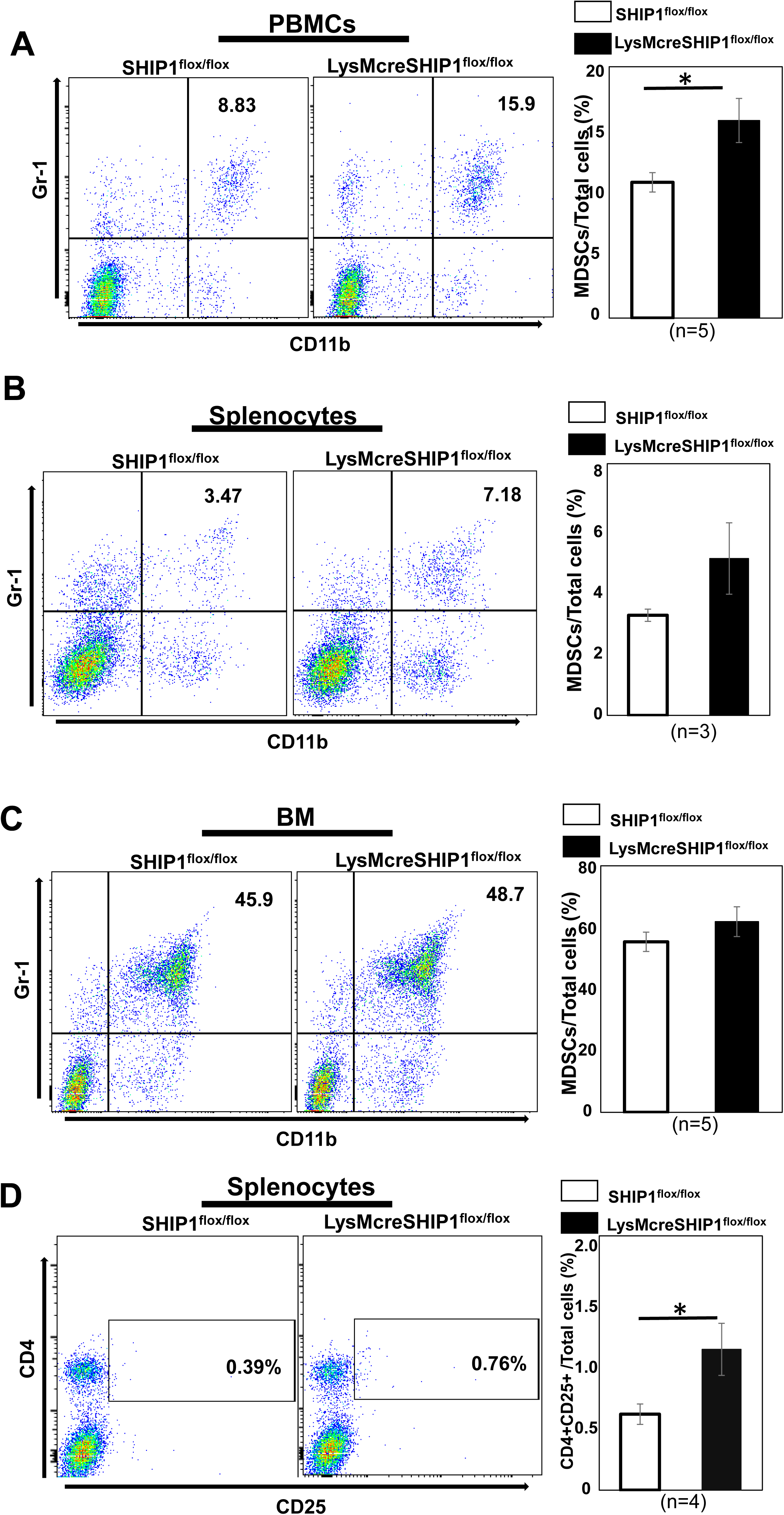
The frequency of regulatory MDSCs and T_reg_ cells in mice with myeloid specific ablation of SHIP1. Single cell suspensions were prepared from peripheral blood (A), spleen (B) and bone marrow (C) of SHIP1^flox/flox^ and LysMcre:SHIP1^flox/flox^ mice. The frequency of CD11b+Gr1+ MDSCs was measured by using a flow cytometer (left panels). The percentage of MDSC population was calculated based on total cell numbers (right panels). (D) The frequency of CD4+CD25+ T_reg_ cells was quantified among total splenocytes. Data are mean ± SEM. *p < 0.05, Student’s t test.

**Figure 2.**
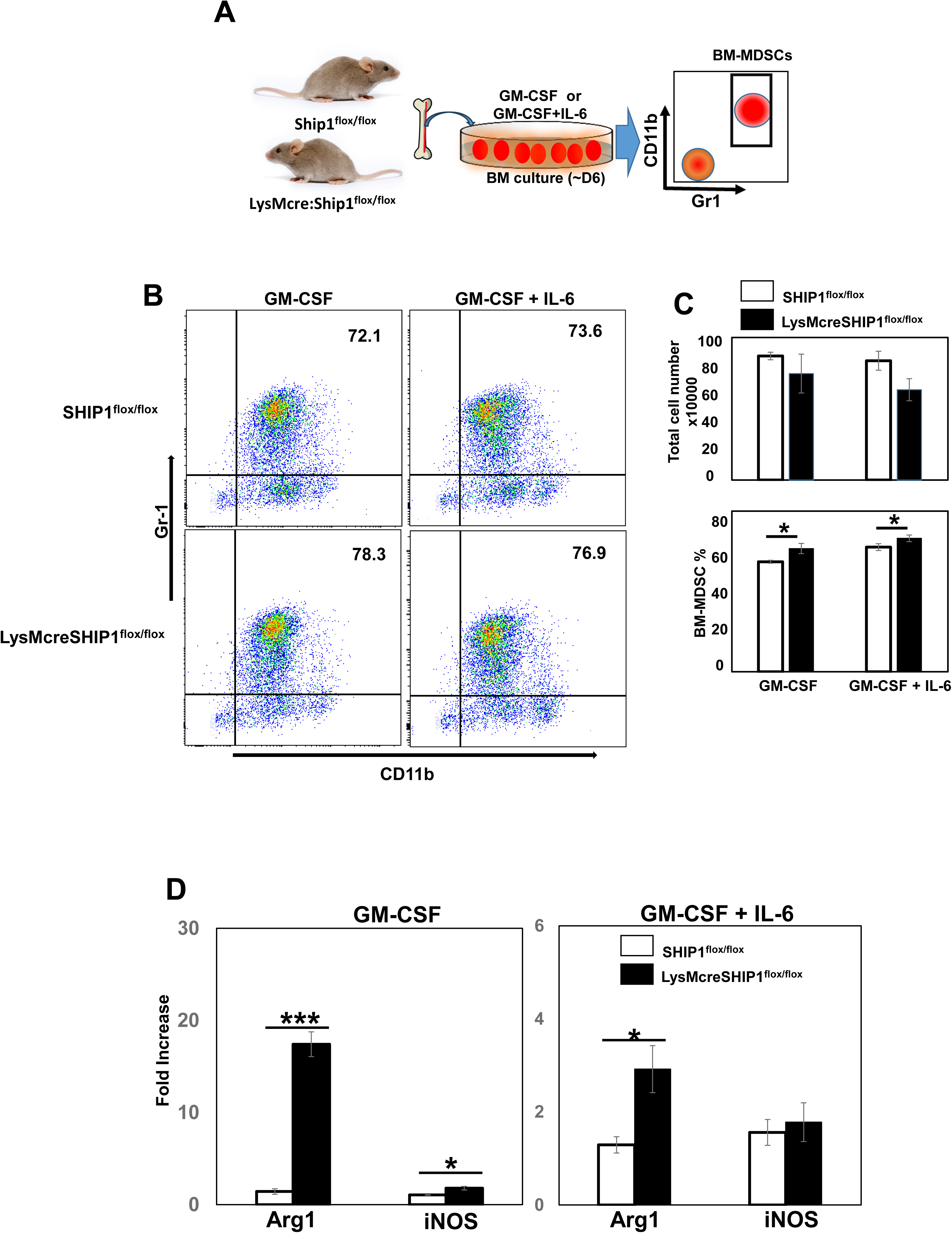
Myeloid specific ablation of SHIP1 promotes the expansion of BM-MDSCs. (A) GM- CSF or GM-CSF+ IL-6 mediated expansion and isolation of BM-MDSCs from BM culture. (B) Flow cytometric analysis of CD11b+Gr1+ MDSCs in BM cultures. Whole BM cells were collected from SHIP1^flox/flox^ and LysMcre:SHIP1^flox/flox^ mice and cultured for 5 days in media containing GM-CSF alone or GM-CSF + IL-6 combination followed by dual staining with anti-mouse CD11b and Gr-1 antibodies. Data are representative of four independent experiments. (C) Total cell numbers were counted by manual cell counting using a hemocytometer (upper panel), and the frequency of BM- MDSCs at Day 5 was expressed as the percentage of CD11b+Gr-1+ cells among total number of cells in the BM culture (lower panel). (D) Total RNA was prepared from BM-MDSCs on Day 4 of SHIP1^flox/flox^ and LysMcre:SHIP1^flox/flox^ BM cultures for gene expression analysis. Following firststrand cDNA synthesis real-time PCR was performed to quantify gene expression levels of the regulatory markers Arg1 and iNOS. GAPDH gene was used as an internal control to validate gene expression. Data are mean ± SEM. ***p<0.001, *p < 0.05, Student’s t test.

### Myeloid specific ablation of SHIP1 enhances the regulatory potential of MDSCs

To further examine the effect of myeloid specific SHIP1 deficiency on MDSC regulatory potential, the expression of functional markers in vitro was analyzed. As shown in Figure 2D, gene expression levels of Arg-1 and iNOS increased significantly in BM-MDSCs from LysMcre:SHIP1^flox/flox^ compared to SHIP1^flox/flox^ control cells. These results suggest that myeloid specific loss of SHIP1 not only increases the frequency of BM-MDSCs but also enhances the regulatory potential of MDSCs. To confirm these results, SHIP1^flox/flox^ BM cells were infected with lentivirus conferring the expression of Cre recombinase and then cultured with GM-CSF and IL-6. Although flox/Cre-induced deletion of SHIP1 did not increase the number of MDSCs (Figure 3A), we observed a significant upregulation of Arg-1 and iNOS gene expression in Cre recombinase-expressing MDSCs compared to control ZsGreen-expressing MDSCs (Figure 3B). Thus, myeloid-specific deletion of SHIP1 augments the immunoregulatory capacity of BM-MDSCs.

**Figure 3.**
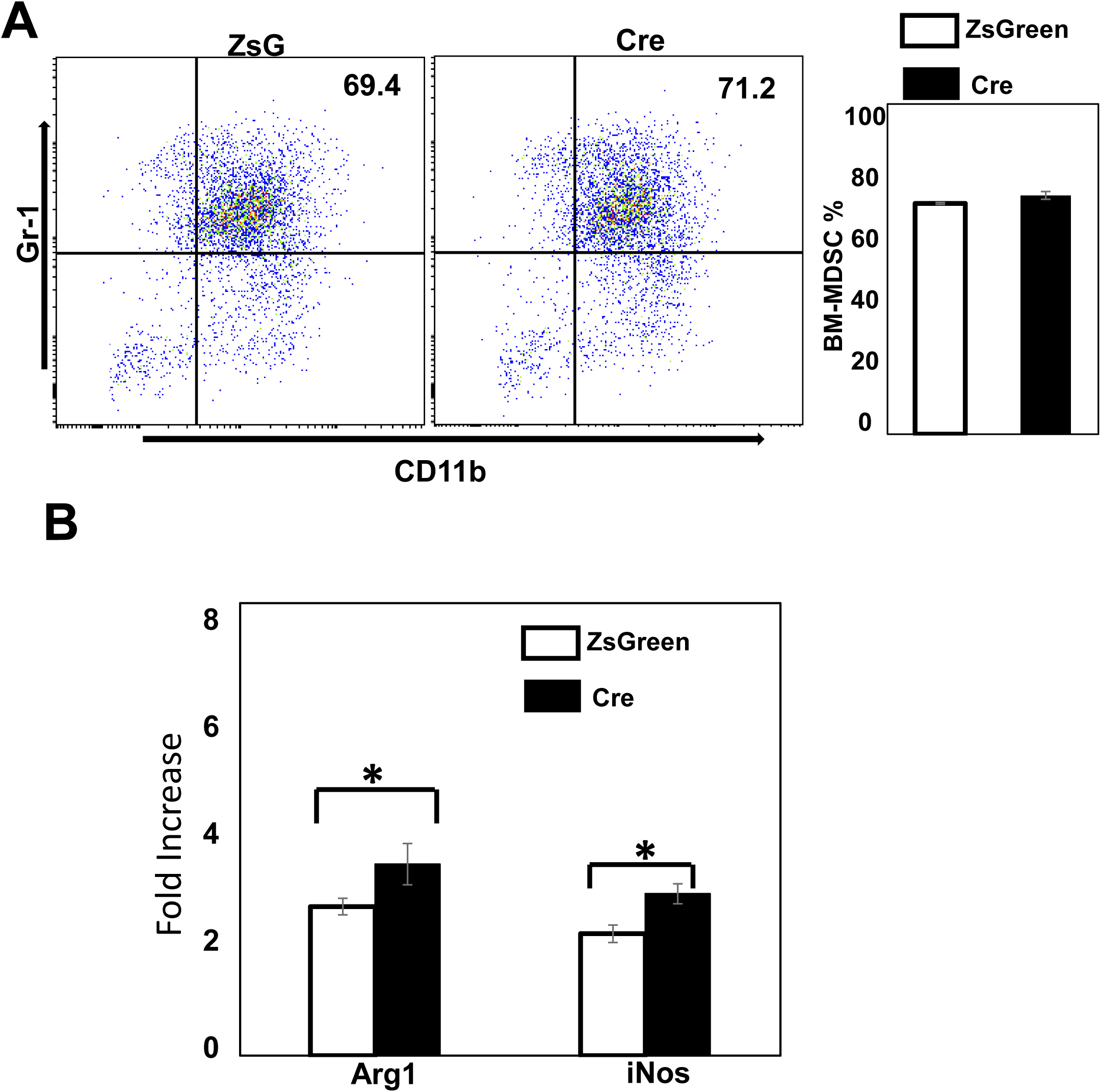
Lentiviral transduction-induced SHIP1 deletion upregulates regulatory gene expression in BM-MDSCs. Whole BM cells were collected from femur and tibia of SHIP1^flox/flox^ mice and cultured in GM-CSF + IL-6 containing media for 4 days. Lentiviruses conferring the expression of fluorescent protein ZsGreen as a control or Cre recombinase for SHIP1 deletion were used to transduce BM cultures through spin-infection at the onset of the experiments. (A) Flow cytometric analysis was conducted to detect the frequency of BM-MDSCs. The results were representative of three independent experiments. (B) Total RNA was prepared from BM-MDSCs at Day 4 of the BM cultures for gene expression analysis. Following first-strand cDNA synthesis real-time PCR was performed to quantify gene expression levels of the regulatory markers Arg1 and iNOS. GAPDH gene was used as an internal control to validate gene expression. Data are mean ± SEM. *p < 0.05, Student’s t test.

### Myeloid specific ablation of SHIP1 augments T cell suppression

To determine if myeloid specific SHIP1 deficiency enhances the immunosuppressive capability of the BM-MDSCs in vitro, we performed T cell proliferation assays in the presence of MDSCs isolated from LysMcre:SHIP1^flox/flox^ or control SHIP1^flox/flox^ BM cultures. We found that BM-MDSCs significantly inhibited the proliferation of both CD4+ and CD8+ T cells in vitro. Importantly, co-culture with LysMcre:SHIP1^flox/flox^ BM-MDSCs more efficiently reduced the percentage of CFSE^low^ T cells than with control BM-MDSCs (Figures 4A, B). Next, we investigated whether myeloid specific loss of SHIP1 can regulate in vivo T cell development in peripheral tissues. We found that the frequency of CD4+ and CD8+ T cells in PBMCs and splenocytes decreased significantly in LysMcre:SHIP1^flox/flox^ mice compared to SHIP1^flox/flox^ control mice (Figures 4C, D), indicating that tissue specific loss of SHIP1 in cells of myeloid lineage can suppresses T cell differentiation and/or proliferation. The reduction of effector T cells in peripheral tissues observed here is like that previously found in global SHIP1 KO mice [38]. These experiments have provided to a limited extent functional validation on enhanced regulatory potential of MDSCs derived from myeloid specific SHIP1 knockout mice. Combined, our results suggest that myeloid specific ablation of SHIP1 augments T cell suppression in vitro and in vivo.

**Figure 4.**
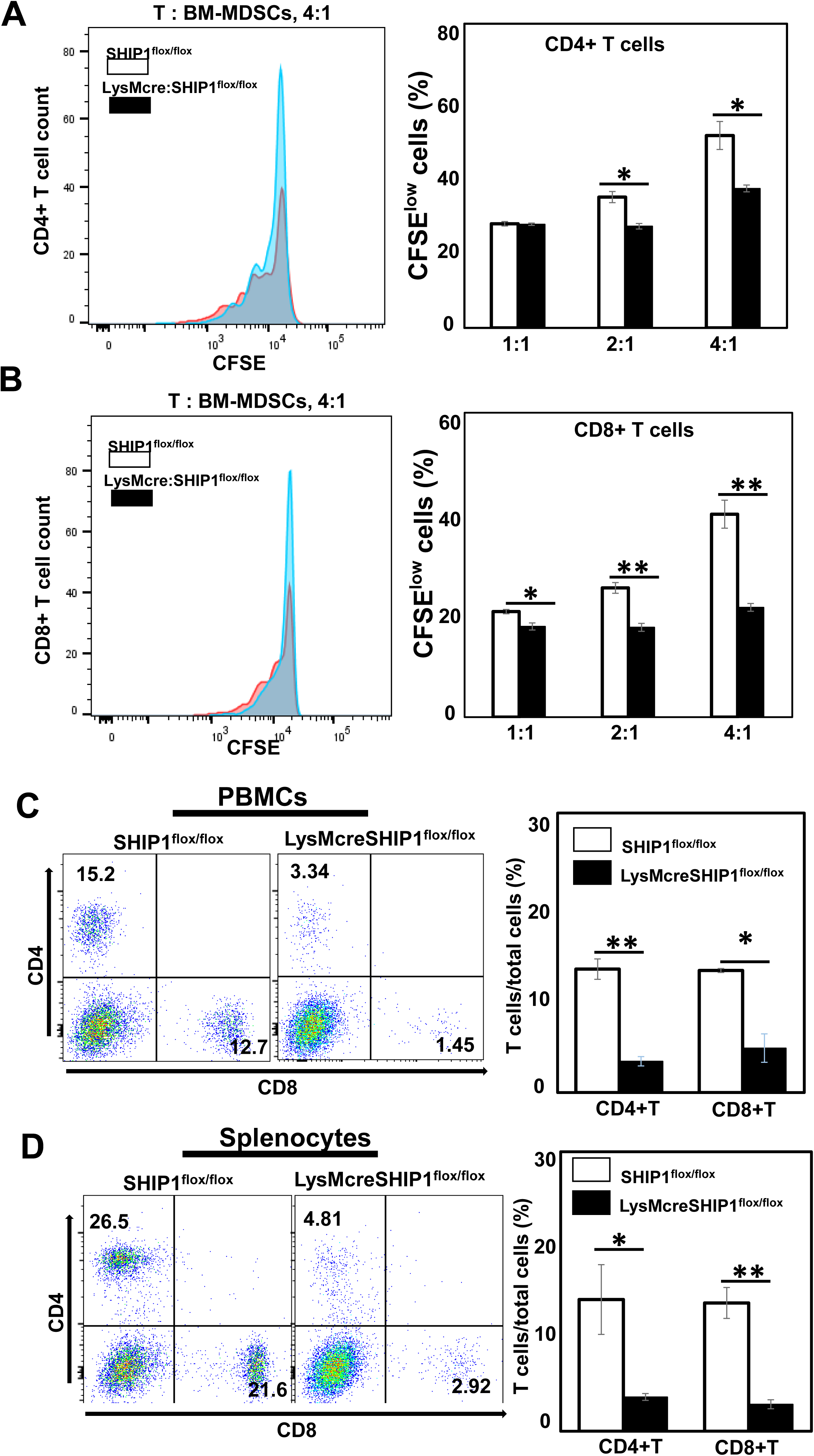
Myeloid specific ablation of SHIP1 augments T cell suppression. Isogenic total T cells were isolated from spleens of SHIP1^flox/flox^ mice and labeled with CFSE before stimulation with Con A. T cells were co-cultured at 1:1, 2:1, and 4:1 ratio with BM-MDSCs generated from GM-CSF containing BM cultures. The percentage of proliferating cells, defined as CFSE^low^, among CD4+ T cells (A) and CD8+ T cells (B) was determined by flow cytometric analysis. (C) and (D) PBMCs and splenocytes were isolated from SHIP1^flox/flox^ and LysMcre:SHIP1^flox/flox^ mice and stained with antiCD4 and CD8 antibodies. The percentage of CD4+ or CD8+ T cells was assessed by flow cytometry. Histograms (left panels) are representative of three independent experiments. Data are mean ± SEM. **p<0.01, *p < 0.05, Student’s t test.

### Myeloid specific ablation of SHIP1 boosts immunoregulatory function of MDSCs in blunting inflammatory arthritis

We next examined the potential of BM-MDSCs as immunomodulatory agents in CIA mouse model. MDSCs expanded and isolated from BM cultures of LysMcre:SHIP1^flox/flox^ mice or SHIP1^flox/flox^ control mice were administered into DBA/1J mice before first and second immunizations to assess the impact on the initiation and severity of inflammatory arthritis (Figure 5A). We photographed mouse paws and performed x-ray imaging at the end of the experiments. Visibly swollen paws and destruction of joint cartilage could be clearly seen in vehicle (PBS) injected CIA mice. In contrast, BM-MDSC injected mice showed nearly no symptom of CIA in photographic and radiographic evaluation (Figure 5B). Furthermore, we found that transplantation of either myeloid SHIP1 KO or control BM-MDSCs significantly suppressed the initiation and progression of CIA in allogeneic DBA/1J mice in comparison with PBS-injected mice (Figure 5C). More importantly, adoptive transfer of myeloid SHIP1 KO MDSCs demonstrated even higher efficacy than control MDSCs in reducing the frequency and severity of CIA development (Figure 5D). These results indicate that myeloid specific ablation of SHIP1 boosts immunoregulatory function of MDSCs in blunting inflammatory arthritis.

**Figure 5.**
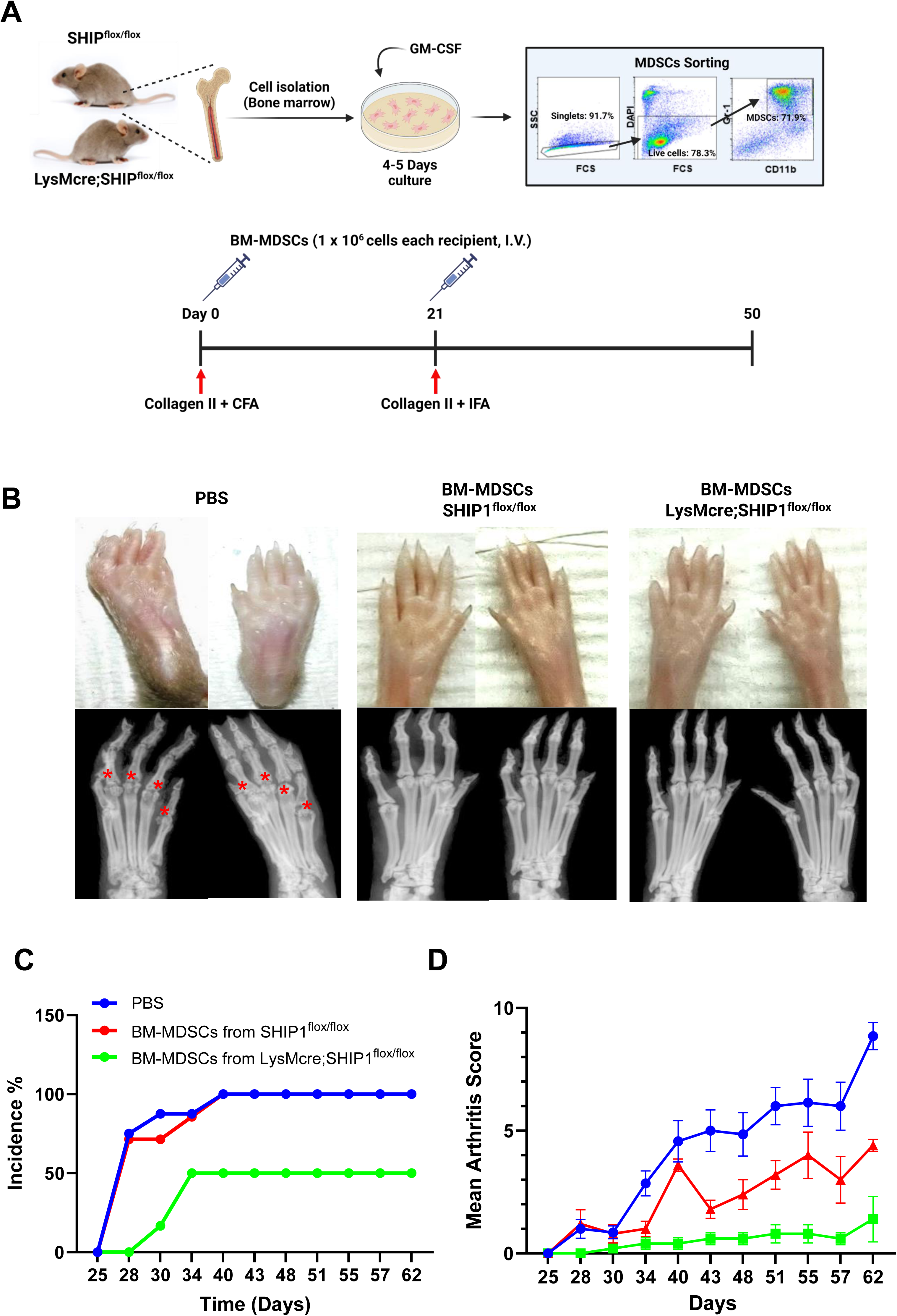
Myeloid specific ablation of SHIP1 boosts immunoregulatory function of MDSCs in blunting inflammatory arthritis in mice. (A) GM-CSF mediated ex vivo expansion and isolation of BM-MDSCs from BM cultures, as well as schematics of the experimental design of an adoptive MDSC transfer in CIA. BM-MDSCs were flow-sorted at Day 4 of BM cultures after staining with anti-CD11b and anti-Gr-1 antibodies. 1 x10^6^ cells were intravenously injected 1 day before each collagen II immunization. (B) Photographic and radiographic images of CIA development. Representative radiographies were from the same front paws as the photographic assessments. Asterisks in red indicate destructive joints of control CIA mice without MDSC injections. The incidence (C) and severity (D) of CIA development in PBS and BM-MDSC recipient mice (n = 5 per group).

### Temporal inhibition of SHIP1 by 3AC in BM culture enhances regulatory functions of BM-MDSCs

In addition, we employed a pharmacological approach by treating bone marrow–derived myeloid-derived suppressor cells (BM-MDSCs) with the SHIP1-specific inhibitor 3AC during their ex vivo culture (Figure 6). Through this experimental strategy, we observed that 3AC-treated BM-MDSCs exhibited a marked increase in the expression of immunoregulatory suppressor genes when compared to control BM-MDSCs (Figure 6B and 6C). To further assess the functional consequences of SHIP1 inhibition, we subsequently evaluated the therapeutic potential of 3AC-conditioned BM-MDSCs in comparison with control MDSCs and vehicle-treated counterparts in the collagen-induced arthritis (CIA) model (Figure 6D and 6E). Consistent with the phenotype observed in genetically SHIP1 deficient MDSCs, the 3AC-conditioned MDSCs demonstrated potent suppressive effects on inflammatory arthritis, which were reflected by a significant attenuation of arthritis incidence and a reduction in clinical symptoms. Collectively, these findings strongly support the conclusion that disruption of SHIP1 signaling in BM-MDSCs enhances their immune-regulatory capacity and further suggest that chemical or genetic targeting of SHIP1 may provide a promising platform for the development of novel MDSC-based cellular therapies for the treatment of RA

**Figure 6.**
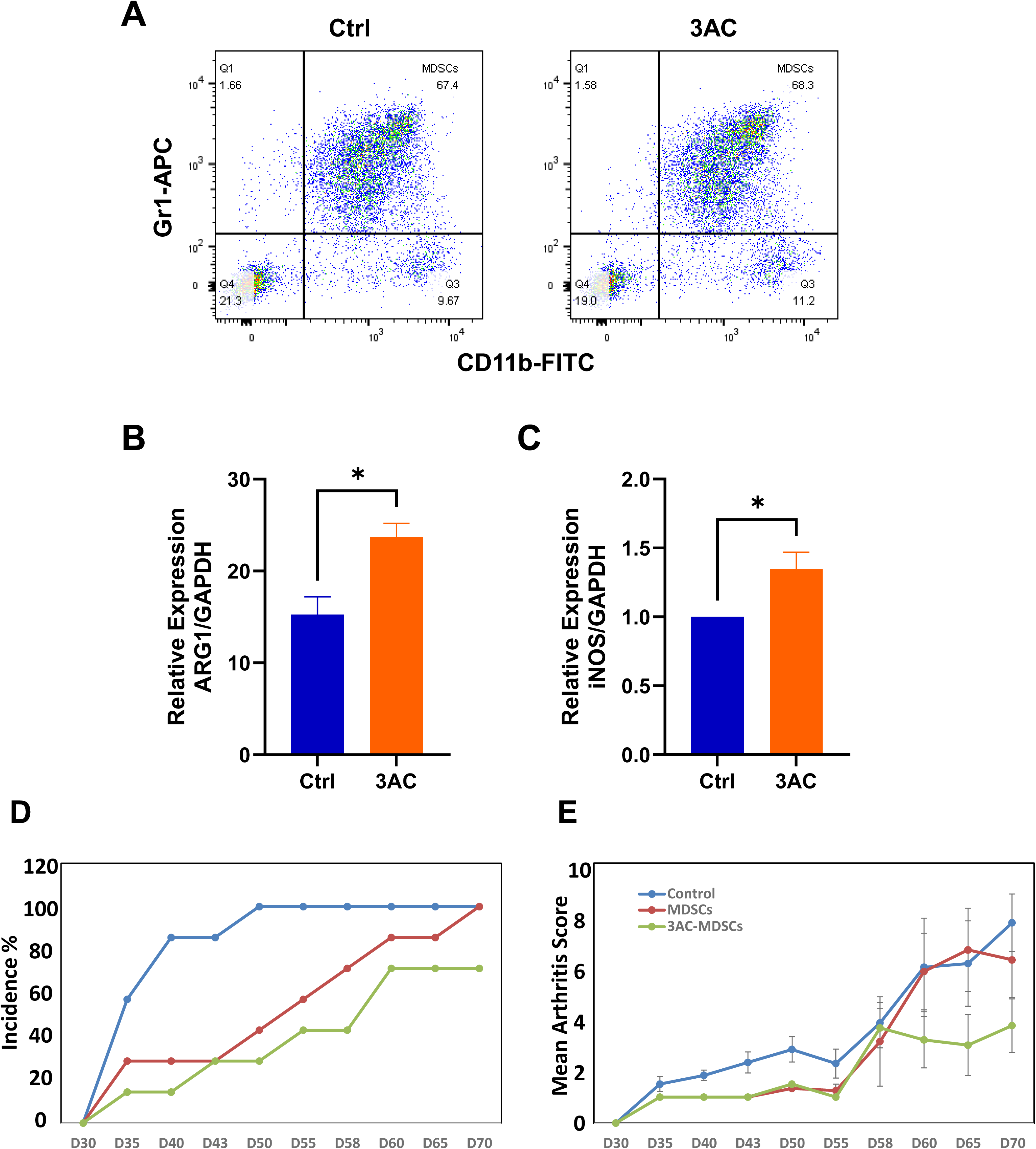
Pharmacological inhibition of SHIP1 increases gene expression of immune regulatory genes in BM-MDSCs and enhances immune regulatory functions of BM-MDSCs. (A) Flow cytometric analysis of CD11b+Gr1+ MDSCs in BM cultures. Whole BM cells were collected from DBA mice and cultured for 5 days in media containing GM-CSF alone (control) and GM-CSF+3AC followed by dual staining with anti-mouse CD11b and Gr-1 antibodies. Data are representative of four independent experiments. Total RNA was prepared from BM-MDSCs on Day 4 for gene expression analysis. Following real-time PCR was performed to quantify gene expression levels of the regulatory markers Arg1(B) and iNOS (C). GAPDH gene was used as an internal control to validate gene expression. BM-MDSCs were flow-sorted at Day 4 of BM cultures after staining with anti-CD11b and anti-Gr-1 antibodies. 1 x10^6^ cells were intravenously injected 1 day before each collagen II immunization. The incidence (D) and severity (E) of CIA development in control and 3AC treated BM-MDSC recipient mice (n = 5 per group). Data are mean ± SEM. ***p<0.001, *p < 0.05, Student’s t test.

## Discussion

A growing number of studies over the past decades have established the key roles of MDSCs in tumor progression and inflammation through suppressive effects on immune cells, mainly by targeting T cell function. The current definition of MDSCs is that they should be of myeloid origin and enriched in mice or patients with cancer or severe disease, display an immature surface phenotype and with the key defining trait being their potent immunosuppressive capacity [31]. However, many questions remain concerning subsets, origin, and function of human MDSCs. In cancer patients, MDSCs are recruited to tumor regions through several chemo-attractants from tumor microenvironment and suppress antitumor responses through inhibitory effects on proliferation and activation of T cells. As such, MDSCs are considered potential therapeutic targets in cancer. On the other hand, MDSCs have also been known as a cellular modulator to regulate autoimmune responses and progression of inflammation. However, findings regarding their immunosuppressive properties have been conflicting. While some studies have identified regulatory properties, others have demonstrated pro-inflammatory involvement through the induction of Th17 lymphocytes [39–41]. Nevertheless, our recent study showed that inhibition of SHIP1 in vivo by a chemical inhibitor 3AC expands MDSCs in vivo and attenuates CIA in mice [24]. Hence, we believe that in our models the potent immunoregulatory capability of MDSCs can be harnessed to mitigate harmful inflammation in RA.

The present study identifies SHIP1 as a central regulator of MDSCs biology and provides novel evidence that myeloid-specific loss of SHIP1 profoundly enhances the immunoregulatory potential of MDSCs in vitro and in vivo. By employing LysMcre:SHIP1^flox/flox^ mice, we demonstrate that this targeted deletion promotes the accumulation of MDSCs in peripheral tissues, enhances their molecular suppressive programs, and confers superior therapeutic efficacy in the CIA model of RA. These findings are further substantiated by pharmacological inhibition of SHIP1 with 3AC, which phenocopied the genetic ablation and highlights SHIP1 as a tractable therapeutic target for ex vivo conditioning of regulatory myeloid cells.

Our results demonstrate that myeloid-specific SHIP1 deletion increases the frequency of MDSCs in spleen and peripheral blood but not in bone marrow (Figure 1), suggesting that SHIP1 primarily regulates peripheral expansion and stability of MDSCs rather than bone marrow progenitor development. Global SHIP1 knockout (KO) mice develop profound hematopoietic abnormalities, including splenomegaly, disrupted lymphopoiesis, and myeloproliferative disease, ultimately leading to reduced lifespan [18, 32, 42]. In contrast, LysMcre:SHIP1^flox/flox^ mice maintain overall immune homeostasis and normal lifespan [35], while still exhibiting a moderate but functionally relevant expansion of MDSCs and regulatory T cells (Tregs), in line with previous studies linking SHIP1 deficiency to enhanced Treg development [17, 43]. Thus, myeloid-specific deletion of SHIP1 uncouples the detrimental hematologic effects of global deficiency from the beneficial regulatory expansion of MDSCs. The observation that bone marrow MDSC frequency was unchanged suggests that SHIP1 exerts its regulatory effect downstream of myelopoiesis, likely by constraining peripheral cues that drive MDSC accumulation. This interpretation is consistent with reports that GM-CSF and IL-6 are critical for MDSC expansion and function [37, 44]. Indeed, our bone marrow cultures revealed that SHIP1-deficient progenitors yielded a significantly higher proportion of MDSCs under GM-CSF or GM-CSF ± IL-6 stimulation (Figure 2), despite comparable total cell numbers to controls. These results indicate that SHIP1 restrains differentiation toward the immature, suppressive MDSC phenotype, favoring instead terminal myeloid maturation.

Myeloid SHIP1 deficiency also enhanced effector programs of MDSCs. Specifically, SHIP1 deficient MDSCs expressed significantly higher levels of Arg-1 and iNOS (Figure 2), two canonical mediators of T cell suppression [8]. Elevated expression of these molecules in SHIP1 deficient MDSCs thus provides a mechanistic explanation for their heightened suppressive activity. These molecular findings translated into superior functional outcomes in vitro, as SHIP1 deficient MDSCs more effectively inhibited both CD4+ and CD8+ T cell proliferation compared with control cells (Figure 4A and B). In vivo, LysMcre:SHIP1^flox/flox^ mice displayed reduced frequencies of effector T cells in spleen and blood (Figure 4C and D), paralleling the in vitro phenotype and underscoring the systemic regulatory effect of SHIP1-deficient MDSCs. This aligns with prior reports showing that global SHIP1 deficiency alters T cell homeostasis [18, 32], but importantly, our data reveal that myeloid-specific SHIP1 ablation alone is sufficient to drive robust suppression of effector T cells.

The CIA experiments highlight the therapeutic potential of SHIP1-deficient MDSCs in autoimmune disease (Figure 5). Adoptive transfer of either SHIP1-deficient or control MDSCs attenuated arthritis incidence and severity, but SHIP1 deficient MDSCs conferred significantly stronger protection, as demonstrated by reduced paw swelling, improved radiographic scores, and lower clinical severity indices. These findings provide direct evidence that SHIP1 deficiency amplifies the disease-modifying function of MDSCs. The CIA model recapitulates key features of human RA, including autoreactive T and B cell responses, synovial inflammation, and joint destruction [45, 46]. While current RA therapies (e.g., TNF-α blockers, JAK inhibitors) primarily target effector immune mechanisms, there is growing recognition that enhancing endogenous regulatory mechanisms could provide durable immune tolerance [47–50]. Our findings suggest that SHIP1-targeted modulation of MDSCs may represent a novel therapeutic avenue that harnesses the body’s own immunoregulatory circuits rather than broadly suppressing immune activity.

A critical translational advance of this study is the demonstration that pharmacological inhibition of SHIP1 with 3AC recapitulates the genetic phenotype. Treatment of bone marrow cultures with 3AC enhanced suppressor gene expression and conferred potent therapeutic efficacy in CIA upon adoptive transfer (Figure 6). This strategy enables transient ex vivo conditioning of MDSCs without the risks associated with systemic genetic ablation of SHIP1. SHIP1 inhibitors such as 3AC have previously been studied in contexts including myeloid malignancy and immune regulation [51, 52]. Our data extend this by providing proof-of-concept that transient pharmacological inhibition can be leveraged to generate regulatory cell products for adoptive therapy. This approach is particularly attractive given the increasing clinical interest in cell-based immunotherapies, where ex vivo manipulation is feasible and avoids systemic off-target toxicity.

SHIP1 negatively regulates PI3K/Akt signaling by dephosphorylating PI(3,4,5)P3 [36]. Loss of SHIP1 thus enhances PI3K/Akt pathway activity, which is known to control myeloid cell survival, metabolism, and differentiation[53]. It is plausible that heightened PI3K/Akt signaling in SHIP1 deficient cells stabilizes an immature myeloid phenotype while activating transcription factors such as STAT3, C/EBPβ, and HIF-1α, all of which are central to MDSC development and function [9]. Therefore, future work should aim to delineate the precise molecular crosstalk between SHIP1-regulated pathways and cytokine-driven transcriptional programs that determine MDSC fate. It will also be important to investigate whether SHIP1 inhibition differentially affects monocytic versus granulocytic MDSC subsets, as these subsets are known to vary in function and therapeutic efficacy [54]. Moreover, given that MDSCs contribute to tumor immune evasion, strategies that enhance MDSC function may carry potential risks in oncology [15, 28]. Balancing therapeutic benefit in autoimmunity with risks in tumor immunity will be an important consideration for clinical translation.

In summary, our findings establish SHIP1 as a pivotal regulator of MDSC biology. Myeloid-specific SHIP1 ablation enhances MDSC frequency, augments their suppressive effector functions, and confers superior protection against autoimmune arthritis. Importantly, transient pharmacological inhibition of SHIP1 reproduces these effects ex vivo, providing a clinically feasible approach to enhance the therapeutic potential of MDSCs. Together, these results suggest that SHIP1-targeted strategies hold promise as a platform for developing MDSC-based therapies for RA and potentially other T cell–mediated autoimmune and inflammatory diseases.

## Data availability statement

The data that support the findings of this study are available from the corresponding author upon reasonable request.

## Ethics statement

All animal experiments were approved by the Brown University Health Institutional Animal Care and Use Committee (IACUC) and carried out in the barrier animal facility at Rhode Island Hospital. All procedures were performed conforming to the NIH Guide for the Care and Use of Laboratory Animals.

## Author contributions

All authors were involved in drafting the manuscript and revising it critically for its scientific content. All authors approved the final version to be published. Dr. Liang has full access to all the data in this study and takes responsibility for the integrity of the data and the accuracy of the data analysis. Study concept and design: EY.S., MJ. C., A.M.R. and O.D.L.; Acquisition of data: EY.S., MJ.C., YE.L., EM.J., A.B., A.M.R. and O.D.L.; Analysis and interpretation of data: EY.S., MJ.C., A.M.R. and O.D.L.

## References

1. Matsuno H, Yudoh K, Katayama R, Nakazawa F, Uzuki M, Sawai T, Yonezawa T, Saeki Y, Panayi GS, Pitzalis C et al: The role of TNF-alpha in the pathogenesis of inflammation and joint destruction in rheumatoid arthritis (RA): a study using a human RA/SCID mouse chimera. Rheumatology (Oxford) 2002, 41(3):329–337.

2. Klareskog L, Catrina AI, Paget S: Rheumatoid arthritis. Lancet 2009, 373(9664):659-672.

3. Boissier MC, Semerano L, Challal S, Saidenberg-Kermanac’h N, Falgarone G: Rheumatoid arthritis: from autoimmunity to synovitis and joint destruction. J Autoimmun 2012, 39(3):222–228.

4. Dadoun S, Zeboulon-Ktorza N, Combescure C, Elhai M, Rozenberg S, Gossec L, Fautrel B: Mortality in rheumatoid arthritis over the last fifty years: systematic review and meta-analysis. Joint Bone Spine 2013, 80(1):29–33.

5. Tu J, Huang W, Zhang W, Mei J, Zhu C: Two Main Cellular Components in Rheumatoid Arthritis: Communication Between T Cells and Fibroblast-Like Synoviocytes in the Joint Synovium. Front Immunol 2022, 13:922111.

6. Kim KW, Kim HR, Kim BM, Cho ML, Lee SH: Th17 cytokines regulate osteoclastogenesis in rheumatoid arthritis. Am J Pathol 2015, 185(11):3011–3024.

7. Wang J, Xue Y, Zhou L: New Classification of Rheumatoid Arthritis Based on Immune Cells and Clinical Characteristics. J Inflamm Res 2024, 17:3293–3305.

8. Gabrilovich DI, Nagaraj S: Myeloid-derived suppressor cells as regulators of the immune system. Nat Rev Immunol 2009, 9(3):162–174.

9. Trikha P, Carson WE, 3rd: Signaling pathways involved in MDSC regulation. Biochim Biophys Acta 2014, 1846(1):55–65.

10. Huang B, Pan PY, Li Q, Sato AI, Levy DE, Bromberg J, Divino CM, Chen SH: Gr-1+CD115+ immature myeloid suppressor cells mediate the development of tumor-induced T regulatory cells and T-cell anergy in tumor-bearing host. Cancer Res 2006, 66(2):1123–1131.

11. Bronte V, Serafini P, Mazzoni A, Segal DM, Zanovello P: L-arginine metabolism in myeloid cells controls T-lymphocyte functions. Trends Immunol 2003, 24(6):302–306.

12. Kusmartsev S, Gabrilovich DI: Inhibition of myeloid cell differentiation in cancer: the role of reactive oxygen species. J Leukoc Biol 2003, 74(2):186–196.

13. Fujii W, Ashihara E, Hirai H, Nagahara H, Kajitani N, Fujioka K, Murakami K, Seno T, Yamamoto A, Ishino H et al: Myeloid-derived suppressor cells play crucial roles in the regulation of mouse collagen-induced arthritis. J Immunol 2013, 191(3):1073–1081.

14. Crook KR, Jin M, Weeks MF, Rampersad RR, Baldi RM, Glekas AS, Shen Y, Esserman DA, Little P, Schwartz TA et al: Myeloid-derived suppressor cells regulate T cell and B cell responses during autoimmune disease. J Leukoc Biol 2015, 97(3):573–582.

15. Fernandes S, Iyer S, Kerr WG: Role of SHIP1 in cancer and mucosal inflammation. Ann N Y Acad Sci 2013, 1280(1):6–10.

16. Ghansah T, Paraiso KH, Highfill S, Desponts C, May S, McIntosh JK, Wang JW, Ninos J, Brayer J, Cheng F et al: Expansion of myeloid suppressor cells in SHIP-deficient mice represses allogeneic T cell responses. J Immunol 2004, 173(12):7324–7330.

17. Kashiwada M, Cattoretti G, McKeag L, Rouse T, Showalter BM, Al-Alem U, Niki M, Pandolfi PP, Field EH, Rothman PB: Downstream of tyrosine kinases-1 and Src homology 2-containing inositol 5’-phosphatase are required for regulation of CD4+CD25+ T cell development. J Immunol 2006, 176(7):3958–3965.

18. Helgason CD, Kalberer CP, Damen JE, Chappel SM, Pineault N, Krystal G, Humphries RK: A dual role for Src homology 2 domain-containing inositol-5-phosphatase (SHIP) in immunity: aberrant development and enhanced function of b lymphocytes in ship -/- mice. J Exp Med 2000, 191(5):781–794.

19. Zhou P, Kitaura H, Teitelbaum SL, Krystal G, Ross FP, Takeshita S: SHIP1 negatively regulates proliferation of osteoclast precursors via Akt-dependent alterations in D-type cyclins and p27. J Immunol 2006, 177(12):8777–8784.

20. Xue H, Hua LM, Guo M, Luo JM: SHIP1 is targeted by miR-155 in acute myeloid leukemia. Oncol Rep 2014, 32(5):2253–2259.

21. Jin HM, Kim TJ, Choi JH, Kim MJ, Cho YN, Nam KI, Kee SJ, Moon JB, Choi SY, Park DJ et al: MicroRNA-155 as a proinflammatory regulator via SHIP-1 down-regulation in acute gouty arthritis. Arthritis Res Ther 2014, 16(2):R88.

22. Yan L, Liang M, Yang T, Ji J, Jose Kumar Sreena GS, Hou X, Cao M, Feng Z: The Immunoregulatory Role of Myeloid-Derived Suppressor Cells in the Pathogenesis of Rheumatoid Arthritis. Front Immunol 2020, 11:568362.

23. So EY, Sun C, Reginato AM, Dubielecka PM, Ouchi T, Liang OD: Loss of lipid phosphatase SHIP1 promotes macrophage differentiation through suppression of dendritic cell differentiation. Cancer Biol Ther 2019, 20(2):201–211.

24. So EY, Sun C, Wu KQ, Dubielecka PM, Reginato AM, Liang OD: Inhibition of lipid phosphatase SHIP1 expands myeloid-derived suppressor cells and attenuates rheumatoid arthritis in mice. Am J Physiol Cell Physiol 2021, 321(3):C569–C584.

25. Bravo A, Kavanaugh A: Bedside to bench: defining the immunopathogenesis of psoriatic arthritis. Nat Rev Rheumatol 2019, 15(11):645–656.

26. Lee SY, Lee SH, Seo HB, Ryu JG, Jung K, Choi JW, Jhun J, Park JS, Kwon JY, Kwok SK et al: Inhibition of IL-17 ameliorates systemic lupus erythematosus in Roquin(san/san) mice through regulating the balance of TFH cells, GC B cells, Treg and Breg. Sci Rep 2019, 9(1):5227.

27. Nishimura K, Saegusa J, Matsuki F, Akashi K, Kageyama G, Morinobu A: Tofacitinib facilitates the expansion of myeloid-derived suppressor cells and ameliorates arthritis in SKG mice. Arthritis Rheumatol 2015, 67(4):893–902.

28. Dufait I, Schwarze JK, Liechtenstein T, Leonard W, Jiang H, Escors D, De Ridder M, Breckpot K: Ex vivo generation of myeloid-derived suppressor cells that model the tumor immunosuppressive environment in colorectal cancer. Oncotarget 2015, 6(14):12369–12382.

29. Ochi K, Morita M, Wilkinson AC, Iwama A, Yamazaki S: Non-conditioned bone marrow chimeric mouse generation using culture-based enrichment of hematopoietic stem and progenitor cells. Nat Commun 2021, 12(1):3568.

30. Yang G, Kramer MG, Fernandez-Ruiz V, Kawa MP, Huang X, Liu Z, Prieto J, Qian C: Development of Endothelial-Specific Single Inducible Lentiviral Vectors for Genetic Engineering of Endothelial Progenitor Cells. Sci Rep 2015, 5:17166.

31. Bronte V, Brandau S, Chen SH, Colombo MP, Frey AB, Greten TF, Mandruzzato S, Murray PJ, Ochoa A, Ostrand-Rosenberg S et al: Recommendations for myeloid-derived suppressor cell nomenclature and characterization standards. Nat Commun 2016, 7:12150.

32. Helgason CD, Damen JE, Rosten P, Grewal R, Sorensen P, Chappel SM, Borowski A, Jirik F, Krystal G, Humphries RK: Targeted disruption of SHIP leads to hemopoietic perturbations, lung pathology, and a shortened life span. Genes Dev 1998, 12(11):1610–1620.

33. Hadidi S, Antignano F, Hughes MR, Wang SK, Snyder K, Sammis GM, Kerr WG, McNagny KM, Zaph C: Myeloid cell-specific expression of Ship1 regulates IL-12 production and immunity to helminth infection. Mucosal Immunol 2012, 5(5):535–543.

34. Collazo MM, Paraiso KH, Park MY, Hazen AL, Kerr WG: Lineage extrinsic and intrinsic control of immunoregulatory cell numbers by SHIP. Eur J Immunol 2012, 42(7):1785–1795.

35. Maxwell MJ, Srivastava N, Park MY, Tsantikos E, Engelman RW, Kerr WG, Hibbs ML: SHIP- 1 deficiency in the myeloid compartment is insufficient to induce myeloid expansion or chronic inflammation. Genes Immun 2014, 15(4):233–240.

36. Liu Q, Sasaki T, Kozieradzki I, Wakeham A, Itie A, Dumont DJ, Penninger JM: SHIP is a negative regulator of growth factor receptor-mediated PKB/Akt activation and myeloid cell survival. Genes Dev 1999, 13(7):786–791.

37. Lee CR, Lee W, Cho SK, Park SG: Characterization of Multiple Cytokine Combinations and TGF-beta on Differentiation and Functions of Myeloid-Derived Suppressor Cells. Int J Mol Sci 2018, 19(3).

38. Anderson CK, Salter AI, Toussaint LE, Reilly EC, Fugere C, Srivastava N, Kerr WG, Brossay L: Role of SHIP1 in Invariant NKT Cell Development and Functions. J Immunol 2015, 195(5):2149–2156.

39. Yi H, Guo C, Yu X, Zuo D, Wang XY: Mouse CD11b+Gr-1+ myeloid cells can promote Th17 cell differentiation and experimental autoimmune encephalomyelitis. J Immunol 2012, 189(9):4295–4304.

40. Zhang H, Wang S, Huang Y, Wang H, Zhao J, Gaskin F, Yang N, Fu SM: Myeloid-derived suppressor cells are proinflammatory and regulate collagen-induced arthritis through manipulating Th17 cell differentiation. Clin Immunol 2015, 157(2):175–186.

41. Guo C, Hu F, Yi H, Feng Z, Li C, Shi L, Li Y, Liu H, Yu X, Wang H et al: Myeloid-derived suppressor cells have a proinflammatory role in the pathogenesis of autoimmune arthritis. Ann Rheum Dis 2016, 75(1):278–285.

42. 42. Kerr WG: Inhibitor and activator: dual functions for SHIP in immunity and cancer. Ann N Y Acad Sci 2011, 1217:1–17.

43. Collazo MM, Wood D, Paraiso KH, Lund E, Engelman RW, Le CT, Stauch D, Kotsch K, Kerr WG: SHIP limits immunoregulatory capacity in the T-cell compartment. Blood 2009, 113(13):2934–2944.

44. Helft J, Bottcher J, Chakravarty P, Zelenay S, Huotari J, Schraml BU, Goubau D, Reis e Sousa C: GM-CSF Mouse Bone Marrow Cultures Comprise a Heterogeneous Population of CD11c(+)MHCII(+) Macrophages and Dendritic Cells. Immunity 2015, 42(6):1197–1211.

45. Brand DD, Latham KA, Rosloniec EF: Collagen-induced arthritis. Nat Protoc 2007, 2(5):1269–1275.

46. Caplazi P, Baca M, Barck K, Carano RA, DeVoss J, Lee WP, Bolon B, Diehl L: Mouse Models of Rheumatoid Arthritis. Vet Pathol 2015, 52(5):819–826.

47. Papagoras C, Drosos AA: Abatacept: a biologic immune modulator for rheumatoid arthritis. Expert Opin Biol Ther 2011, 11(8):1113–1129.

48. O’Shea JJ, Schwartz DM, Villarino AV, Gadina M, McInnes IB, Laurence A: The JAK-STAT pathway: impact on human disease and therapeutic intervention. Annu Rev Med 2015, 66:311–328.

49. Aaltonen KJ, Virkki LM, Malmivaara A, Konttinen YT, Nordstrom DC, Blom M: Systematic review and meta-analysis of the efficacy and safety of existing TNF blocking agents in treatment of rheumatoid arthritis. PLoS One 2012, 7(1):e30275.

50. Schioppo T, Ingegnoli F: Current perspective on rituximab in rheumatic diseases. Drug Des Devel Ther 2017, 11:2891–2904.

51. Brooks R, Fuhler GM, Iyer S, Smith MJ, Park MY, Paraiso KH, Engelman RW, Kerr WG: SHIP1 inhibition increases immunoregulatory capacity and triggers apoptosis of hematopoietic cancer cells. J Immunol 2010, 184(7):3582–3589.

52. Saz-Leal P, Del Fresno C, Brandi P, Martinez-Cano S, Dungan OM, Chisholm JD, Kerr WG, Sancho D: Targeting SHIP-1 in Myeloid Cells Enhances Trained Immunity and Boosts Response to Infection. Cell Rep 2018, 25(5):1118–1126.

53. Yu JS, Cui W: Proliferation, survival and metabolism: the role of PI3K/AKT/mTOR signalling in pluripotency and cell fate determination. Development 2016, 143(17):3050–3060.

54. Ferrer G, Jung B, Chiu PY, Aslam R, Palacios F, Mazzarello AN, Vergani S, Bagnara D, Chen SS, Yancopoulos S et al: Myeloid-derived suppressor cell subtypes differentially influence T-cell function, T-helper subset differentiation, and clinical course in CLL. Leukemia 2021, 35(11):3163–3175.

