## Supplemental Figure 1 for "Myeloid Specific Ablation of SHIP1 Boosts ex vivo Expansion and Regulatory Function of Myeloid-Derived Suppressor Cells in Inflammatory Arthritis"

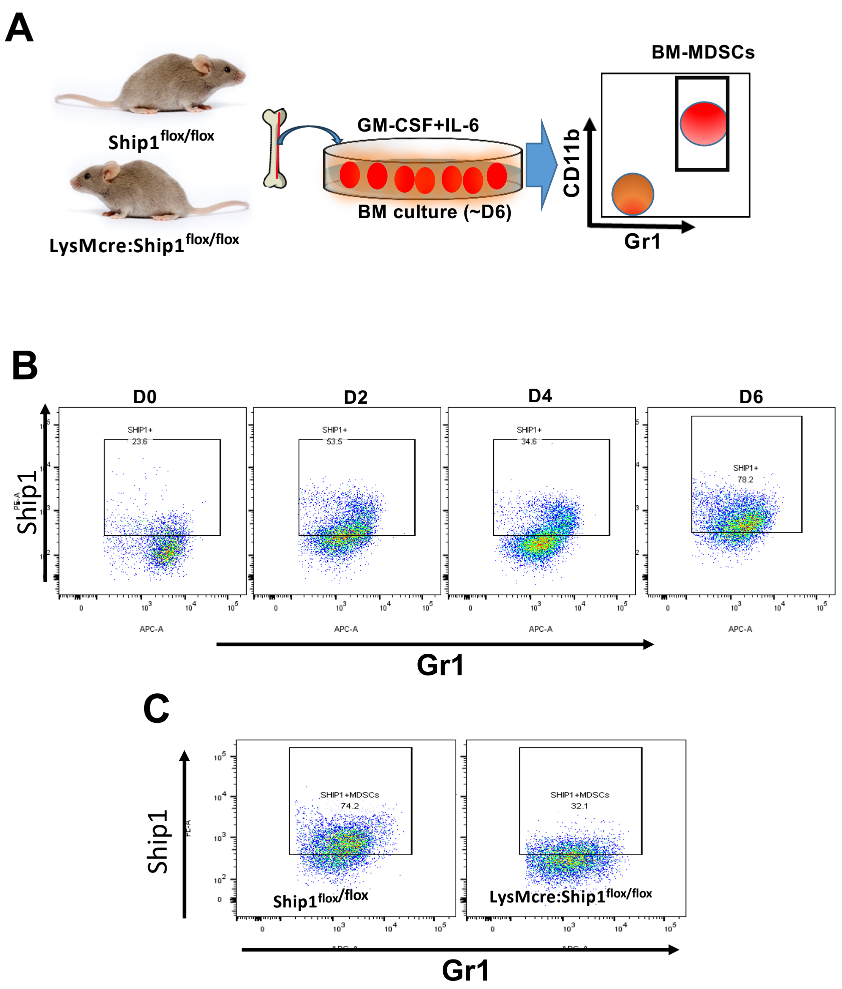


BM cells were isolated from SHIP1$^{flox/flox}$ (B and C), and LyMCre;Ship1$^{flox/flox}$ (C) mice as described in Materials and Methods (A). Cells were cultured in RPMI 1640 medium supplemented with GM-CSF (10 ng/ml) and IL-6 (10 ng/ml) for 6 days. BM-MDSCs were harvested at the indicated time points and stained with APC–anti-mouse Gr1 and Alexa Fluor 488–anti-mouse CD11b antibodies prior to intracellular staining. Intracellular staining for SHIP1 was performed using Fixation Buffer and Perm/Wash Buffer (BioLegend, CA), according to the manufacturer's instructions. PE–anti-SHIP1 antibody was used to determine SHIP1 protein expression in BM-MDSCs. During BM culture, SHIP1 expression increased in a time-dependent manner and reached its maximum level at day 6 (B). BM-MDSCs from LysMcre;Ship1$^{flox/flox}$ mice showed significantly reduced expression (C). Data are representative of three independent experiments.
